# Nuclear partitioning by surface condensation on metaphase chromatids

**DOI:** 10.1101/2024.11.13.622586

**Authors:** Vasanthanarayan Murugesan, Thomas Quail, Maria Elsner, Jan Brugués

## Abstract

Protein partitioning between the nucleus and the cytoplasm is a defining feature of eukaryotic cells, and it is thought to be primarily regulated by transport across the nuclear envelope^1–3^. However, during early vertebrate development, ensuring accurate nuclear composition is challenging due to dramatic changes in cell size and rapid cell division^4–7^. Here, we show that the embryonic linker histone H1.8 is rapidly partitioned into the nucleus through a nuclear import-independent mechanism. In the cytoplasm of *Xenopus laevis* egg extracts, prior to nuclear assembly, H1.8 condenses on the surface of chromatids as droplets that partially wet them. Simultaneously, H1.8 also forms condensates in the cytoplasm that buffer the condensate nucleation rate on the chromatid surface, resulting in a nuclear partitioning that is independent of the DNA-to-cytoplasm ratio. To show the generality of the surface condensation mechanism, we extended our study to Nucleophosmin 1 (NPM1) in human cell culture. Similar to H1.8, NPM1 forms a layer around chromatids that breaks into droplets that subsequently are incorporated into the nucleus. Our findings show that the properties of condensate wetting and nucleation on chromatid surfaces provide an alternative and robust biophysical mechanism to regulate nuclear composition.

## Main Text

Nuclear composition and accurate protein partitioning between the nucleus and the cytoplasm is thought to be primarily regulated by the gradual import of proteins across the nuclear envelope. The nuclear amounts of many imported proteins vary significantly during embryonic development as a consequence of dramatic changes in the nuclear-to-cytoplasmic ratio^4–6,8–12^. These changes in amounts enable the embryo to activate transcription, cell motility, and heterochromatin formation. However, the amounts of some key structural proteins, such as the linker histone, is maintained to ensure proper DNA packaging and function^13,14^. It is unclear how nuclear import can be compatible with these two opposed dependencies of nuclear amounts. Here, we unravel an alternative mechanism for nuclear entry that is independent of nuclear import. This alternative mechanism is based on the nucleation of condensates on chromatid surfaces prior to nuclear assembly and can lead to nuclear amounts that are insensitive to the nuclear-to-cytoplasmic ratio.

### An import-independent mechanism drives H1.8 partitioning in the nucleus

We used nuclei assembled in *X. laevis* cycling egg extracts as an assay to investigate how protein concentration in the nucleus^15–17^ is regulated during embryonic development. Adding demembranated frog sperm isolated from male frogs to extracts generates pronuclei that are functional and undergo DNA replication. In interphase, these nuclei import proteins through nuclear pore complexes and increase in volume. Using GST-mCherry-NLS as a proxy for bulk import, we observed a monotonic increase in the mean intensity as the nucleus grew, as previously reported for a continuously imported component^8,18^. This increase in mean intensity continued until nuclear disassembly during prophase at which point the signal vanished (Fig.1 a, b). This behavior is typical of a protein that undergoes rapid import.

**Fig. 1.**
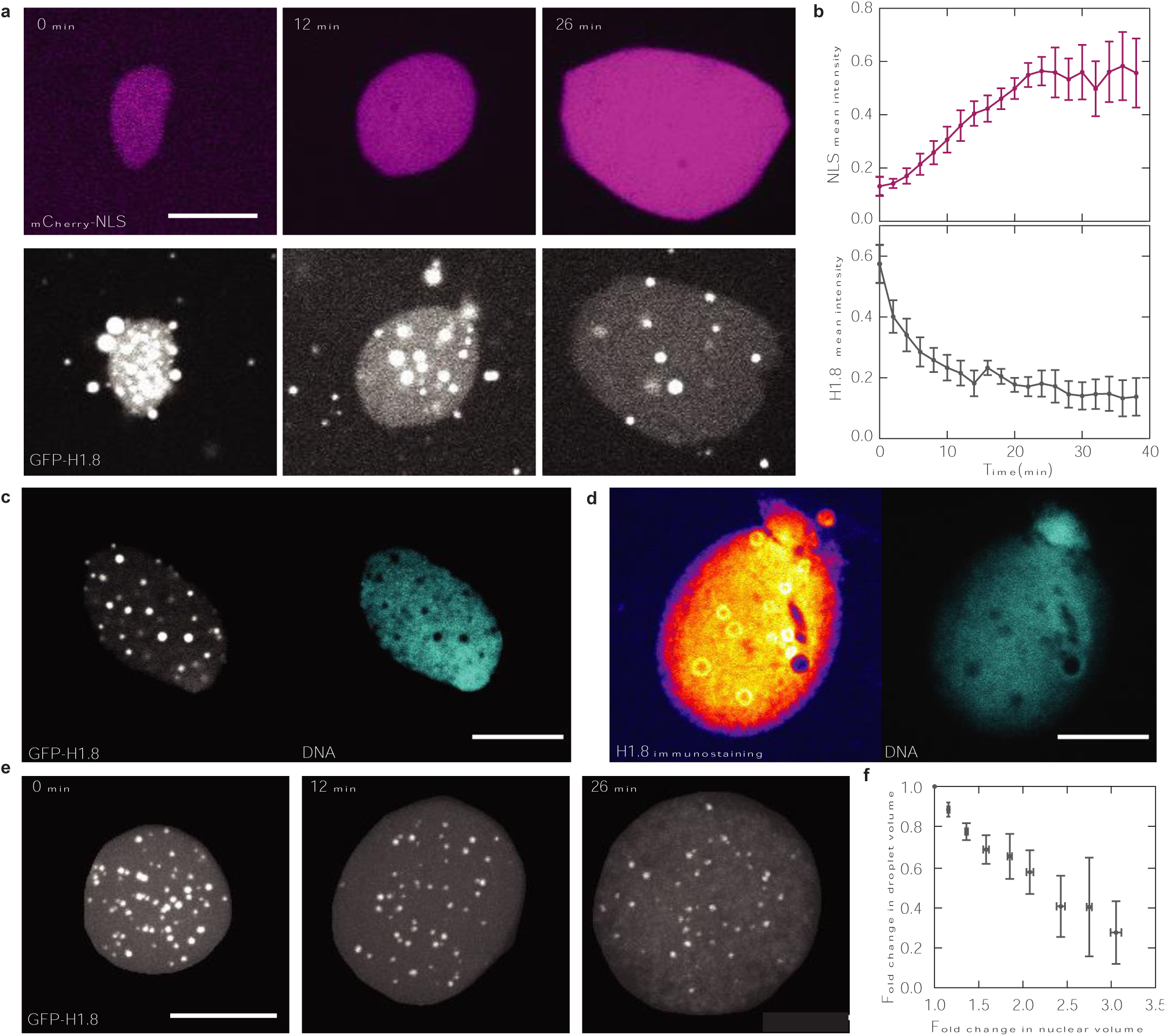
H1.8 is incorporated into the nucleus by a mechanism other than import. a, Representative time lapse images of nuclear growth imaged with mCherry-NLS (concentration 1 µM) and GFP-H1.8 (concentration 2.8 µM). b, Quantification of nuclear mean intensity (total intensity/area) shows that the mCherry-NLS concentration increases with time while GFP-H1.8 concentration decreases (mean ± s.e.m, n = 55 nuclei from 5 interphases). c, Representative image of a nucleus with GFP-H1.8 and DNA shows that DNA is absent in GFP-H1.8 nuclear condensates. d, Representative immunofluorescence image stained against H1.8 and DNA show that H1.8 condensates exclude DNA. e, Representative maximum intensity projection of a nucleus with GFP-H1.8 shows that GFP-H1.8 condensates dissolve as the nucleus grows. f, Total condensate volume decreases with increasing nuclear volume (mean ± s.e.m, n=6). All scale bars, 10 µm.

The embryonic linker histone H1.8 (also known as H1OO, H1foo, H1M, and B4) is the major linker histone variant present in vertebrate oocyte and early embryos and has been shown to affect chromatin structure and mitotic chromatid architecture in a concentration dependent manner^19–23^. H1.8 is expected to follow a similar import dynamic to NLS as it is one of the earliest proteins to partition within the nucleus during early development^5^.

Surprisingly, GFP-H1.8 showed the opposite behavior, with its nuclear concentration peaking at the beginning of interphase and monotonically decreasing as nuclei grew in volume (Fig. 1a, b). Additionally, while H1.8 was bound to chromatin, we also noticed the presence of distinct foci both in the nucleus and the cytoplasm (Fig. 1a). H1.8 foci exhibited characteristic features of liquid-liquid phase separation (Extended Fig. 1) and excluded chromatin in the nucleus (Fig. 1c, d, and Extended Fig. 2), showing an excess of nuclear H1.8 that does not bind to chromatin. The volume of these condensates decreased with increasing nuclear volume (Fig. 1e, f and Supplementary Video 1), consistent with a decreasing concentration of H1.8 in the nucleus due to minimal import. This decrease in condensate volume suggests that H1.8 condensates could act as stores of excess protein that may be readily released for processes such as rapid replication within the nucleus, given that H1.8 is not enriched during interphase. These results are consistent with H1.8 not binding to importin β^24^ and its concentration is buffered during early embryogenesis^14^. In contrast, a somatic variant of linker histone did not phase separate but instead it was imported^24^ and dramatically compacted chromatin^25^ within the nucleus (Extended Fig. 3). Disruption of the disordered C-terminus of H1.8 resulted in the abolishment of phase separation and the formation of nuclear condensates. This indicates that condensation is the driving force behind the creation of nuclear stores, and suggests that distinct charge distributions in the C-termini between somatic and embryonic variants may account for their disparate condensation behaviours (Extended Fig. 4). Altogether, these data suggest that most embryonic H1.8 in the nucleus is not added during nuclear growth and consequently its partitioning in the nucleus depends on mechanisms other than import.

### H1.8 condenses on the chromatid surface

A plausible mechanism to enrich H1.8 in the nucleus is through chromatin binding before nuclear formation because of its known affinity for mitotic chromatids^24,26^. However, the role of its binding affinity to chromatin and its impact on nuclear partitioning has not been assessed. Additionally, how H1.8 condensates would form within the nucleus is unclear for two reasons. First, nuclear condensates typically form beyond a threshold concentration after the import of its constituent proteins^27,28^. Second, even if the protein binds to chromatids, it is unclear how stoichiometric binding could lead to the excess protein in the nucleus needed to form condensates. To investigate the enrichment of H1.8 in the nucleus, we imaged H1.8 dynamics during the transition from chromatid decondensation to nuclear formation in extract. During this process, we observed the emergence of small distinct foci on the chromatin surface that increased in size before mCherry-NLS import (Fig. 2a Supplementary Video 2). Immunostaining of native H1.8 also suggested the presence of H1.8 foci on chromatid surface at physiological concentration (Fig. 2b). These results support the hypothesis that H1.8 is enriched in the nucleus by nucleation of H1.8 condensates on the chromatid surface.

**Fig. 2.**
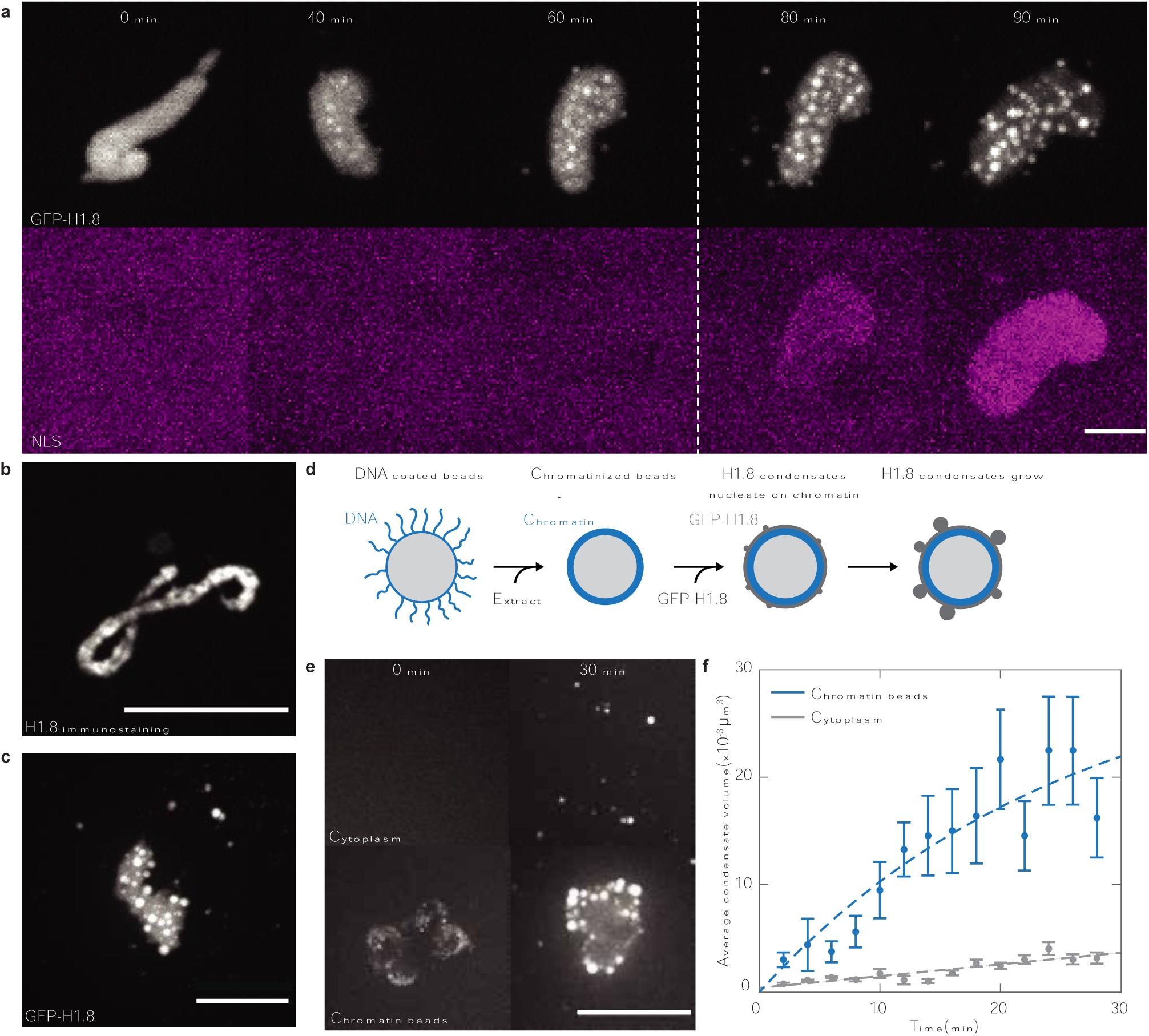
H1.8 condenses on the surfaces of chromatids prior to nuclear assembly. a, Time lapse imaging of nuclear formation from sperm chromatid shows that H1.8 condensates form before the import of NLS protein (marked by a dashed white line). b, Representative immunofluorescence image of chromatid stained with H1.8 reveals that H1.8 form foci on the chromatid surface. c, Chromatid in metaphase arrested extract with GFP-H1.8 (concentration 2.8 µM) shows that H1.8 form condensates on the surface of chromatid. d, Schematic of chromatinized DNA beads nucleating H1.8 condensates. e, Representative images of H1.8 condensate formation in the cytoplasm and on chromatinized beads show that condensates have a preference to nucleate on the chromatin surface. f, Quantification of H1.8 condensate dynamics on the chromatin surface (yellow) and bulk cytoplasm (blue), mean ± s.e.m, n=24 for chromatids, and n=70 for cytoplasm. Dashed lines are fits for condensate volume growth that is proportional to the available chromatin surface (yellow) and a constant bulk rate (blue). All scale bars, 5 µm.

To test this hypothesis—and to exclude contributions from other surface processes in nucleogenesis such as LEM2 binding^29^, and the formation of the nuclear enevelope^30^—we visualized H1.8 localization on chromatin in CSF-arrested extracts. In CSF-arrested extracts, the cell cycle is arrested in metaphase unless triggered into interphase by the addition of calcium to the extracts^15^. In this condition, we observed droplets that partially wetted the surface of the chromatids (Fig. 2c). These experiments suggest that chromatids alone can promote H1.8 condensate formation.

To test whether the chromatin surface was sufficient to nucleate those condensates, we added DNA-coated beads into CSF-arrested extracts, leading to chromatin assembly on the bead surfaces (Fig. 2d). In this condition, we confirmed that H1.8 condensates formed on the surface and grew much faster than condensates formed in the cytoplasm (Fig. 2e, Supplementary Video 3). The simple geometry of the beads allowed us to measure an average contact angle of 119°± 1.4° for the condensates formed on the beads (Extended Fig. 5). Quantification of the growth kinetics in the cytoplasm (bulk) and DNA-coated beads (surface) showed that that kinetics of surface nucleation was 10-fold larger compared with bulk nucleation (Fig. 2f). Surface nucleation was consistent with a rate proportional to the available chromatin surface^31^. Altogether, these data show that enrichment of H1.8 in the nucleus is mediated by surface nucleation of condensates that wet the chromatids during metaphase and are subsequently incorporated into the nucleus. This mechanism is independent of nuclear import and depends on the nucleation rate on chromatids.

### Cytoplasmic condensates buffer surface nucleation on chromatids

The rate of surface nucleation on mitotic chromatids should depend on the soluble concentration of protein in the cytoplasm^28,31^. The soluble concentration can be affected by changes in the DNA-to-cytoplasmic ratio (methods), a crucial regulator of embryonic processes^5–7,32^. However, since H1.8 formed condensates in the cytoplasm, we expected its soluble concentration, and thus the nucleation rate on chromatids, to be buffered to changes in total amount of protein. This is because the formation of biomolecular condensates takes place at a critical concentration above which the concentration of the non-condensed phase is buffered^33,34^.

To test this idea, we used an oil-extract emulsion to encapsulate cytoplasmic extract with sperm DNA in droplets of different volumes^35^. This approach allowed us to vary the total amount of protein and DNA-to-cytoplasmic ratio (Extended Fig. 6) and then quantify surface nucleation of H1.8 in the presence and absence of cytoplasmic condensates. We first exploited the fact that the formation of cytoplasmic condensates is slow (see Fig 2 e, f) to test the dependence of surface nucleation on the total amount of cytoplasmic protein. To this end, we added sperm DNA into CSF-arrested extract at 4 °C, which prevents the formation of H1.8 cytoplasmic condensates. We then encapsulated these reactions in oil at room temperature (Fig. 3a). After 30 min of incubation, the volume of the condensed phase scaled with the cytoplasmic volume (Fig.3 b, c). Thus, in the absence of cytoplasmic condensates, H1.8 condensation on chromatids is consistent with surface nucleation that depends on a limited pool of cytoplasmic protein.

**Fig. 3.**
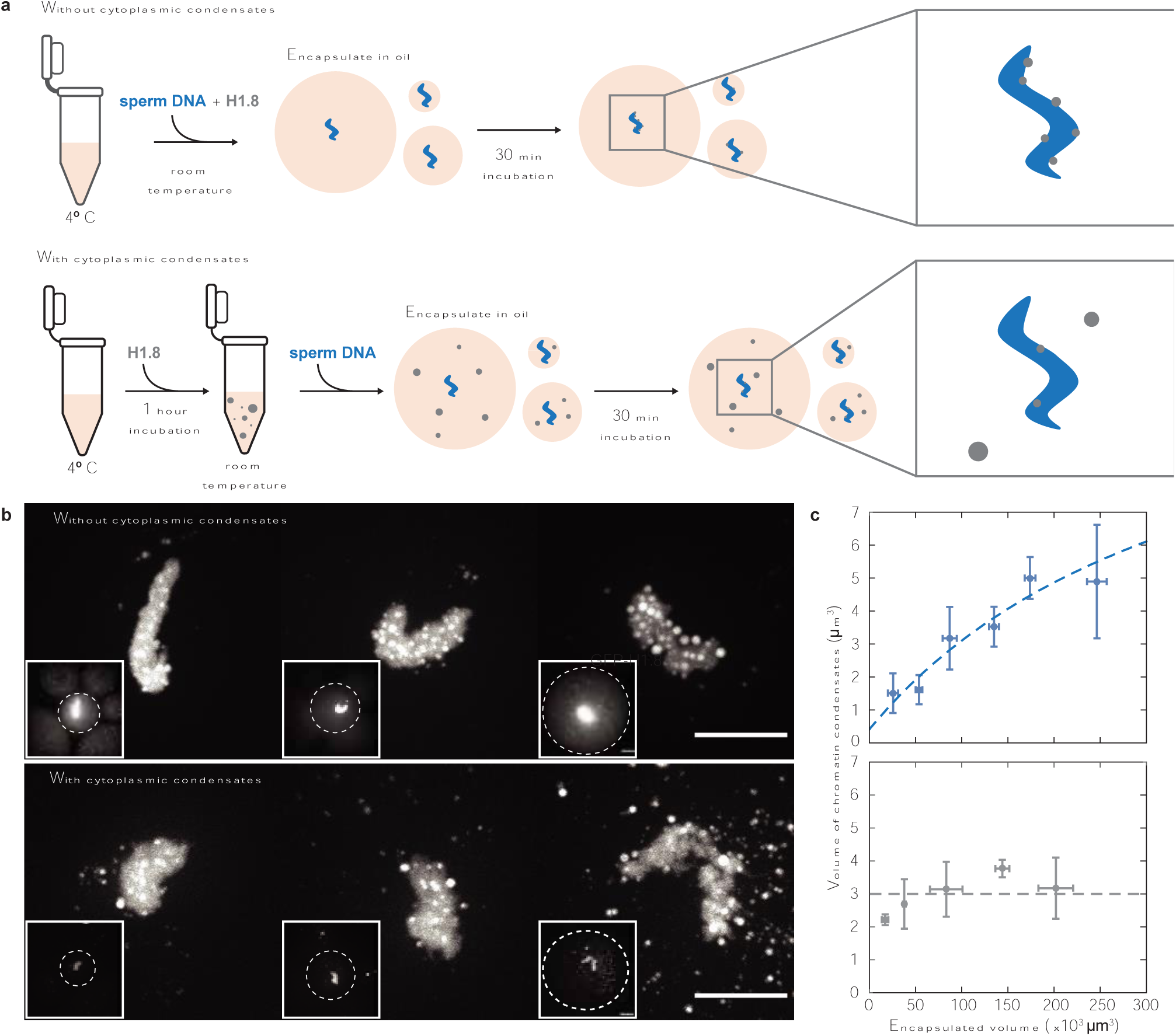
Cytoplasmic condensates buffer surface nucleation on chromatids. a, Schematics showing the experimental assay to study the effect of H1.8 nucleation across different cytoplasmic volumes in the absence (top) and presence (bottom) of cytoplasmic H1.8 condensates. b, Maximum intensity projections of chromatids wetted by H1.8 surface condensates in increasing cytoplasmic volumes (left to right). Inset shows the volume of encapsulated extract (dashed white line) and the dynamic range is changed to highlight the cytoplasmic condensates. The total amount of surface condensates scales with cytoplasmic volume in the absence of cytoplasmic condensates (top) but remains constant when cytoplasmic condensates are present (bottom). Scale bar 10 µm. c, Quantification of surface condensates versus cytoplasmic volume fitted with the scaling model (described in methods) in the absence of cytoplasmic volume (top, n=39) and fitted with a constant in the presence of cytoplasmic condensates (bottom, n=34).

Next, we incubated CSF extract for 1 hour at room temperature to allow for the formation of H1.8 cytoplasmic condensates prior to adding sperm DNA. After 1 hour, we added sperm DNA and encapsulated the extract in oil (Fig. 3a). In this situation, the dilute concentration is independent of the volume of the cytoplasmic condensed phase, and thus the encapsulation volume. Consistent with buffering of the surface condensation rate, the volume of surface condensates after 30 min was independent of the extract encapsulation volume (Fig. 3 b, c). Taken together, our data show that cytoplasmic condensation leads to a rate of surface nucleation on mitotic chromatids that is independent of the nuclear-to-cytoplasmic ratio.

### Nucleolar protein NPM1 partitions into the nucleus through surface condensation on chromatids in human cell lines

Our data suggests that surface condensation on mitotic chromatids may be a general mechanism to partition proteins into the nucleus independently of nuclear import in early embryogenesis. To test this mechanism in other contexts, we investigated how the nucleolar protein NPM1 enters the nucleus in transformed human cell lines. Previous research has shown that the presence of a thin layer of NPM1 on the mitotic chromatids is necessary for its proper nuclear localization^36,37^, suggesting that surface condensation may drive its localization into the nucleus. However, due to phosphorylation, NPM1 is diffuse during metaphase^38^ and hence it is unclear whether condensation plays a role in its nuclear localization.

We imaged the localization of native NPM1, tagged with mNeonGreen using CRISPR-Cas9^33^, as the cell transitions from metaphase to interphase. In metaphase, NPM1 is mostly diffusive in the cytoplasm except for the thin layer present on the chromatid surface. As it enters anaphase, NPM1 is enriched on the chromatid surface, and as anaphase progresses it is broken down into smaller droplets that wet the surface (Fig. 4 a, b and Supplementary Video 4), similar to H1.8. As chromosomes cluster and enter telophase, NPM1 condensates localize within the chromatin matrix before the formation of a nuclear envolope (Fig. 4 a,b). Upon quantification of NPM1 intensity on the chromatin surface and within the chromatin matrix over time, we observed an increase in the concentration of NPM1 inside the chromatin matrix (Fig. 4 c). This increase coincided with a decrease in NPM1 concentration on the chromatin surface, indicating that NPM1, which initially condensed on the chromatin surface, subsequently entered the chromatin matrix. Similar to H1.8, NPM1 also forms condensates in the cytoplasm. Consistent with the chromatin surface promoting condensation, we observed that NPM1 first condenses on the chromatid surface before bulk condensation in the cytoplasm (Extended Fig. 7 a), suggesting a possible role on regulating the total amount of NPM1 entering the nucleus (Extended Fig. 7 b). In summary, these findings indicate that the principles of surface condensation driving nuclear partitioning that we uncovered using H1.8 can be also extended to other *in vivo* systems.

**Fig. 4.**
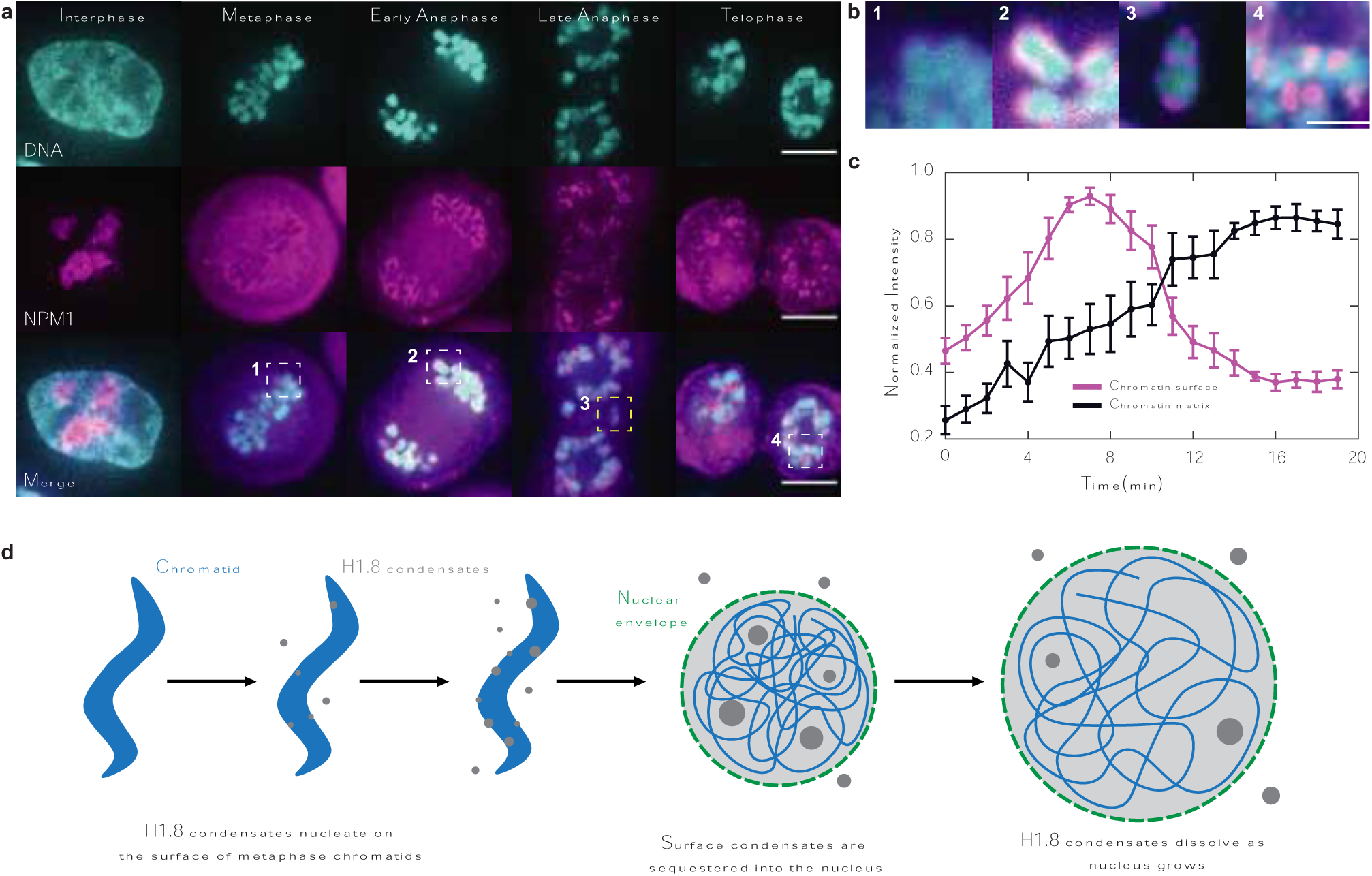
Surface condensation on chromatids facilitates the nuclear partitioning of the nucleolar protein NPM1. a, Representative images of NPM1 localisation in different stages of cell cycle in HCT116 cell lines. Scale bar 5 µm. b, Magnified images of the white dashed box in figure a to highlight that NPM1 translocates from the surface to the chromatin matrix. Scale bar 2 µm c, Quantification of mean intensity of NPM1 on the surface (magenta) and within the chromatin matrix (black) as cells transitions from metaphase to interphase. (mean ± s.e.m, n=7). d, Model of H1.8 partitioning into the nucleus. Condensates of H1.8 are nucleated on the surface of metaphase chromatids, and in the cytoplasm, and increase in size over time. As the cell cycle progresses into interphase, the surface condensates enter the nucleus and acts as reservoirs of non-DNA bound protein that dissolve as the nucleus grows.

## Discussion

Our study reveals that surface condensation and wetting of proteins on chromatids provide a mechanism to regulate nuclear composition (Fig. 4 d). This mechanism results in the immediate availability of protein reservoirs after nuclear envelope formation, as opposed to the gradual concentration of components via nuclear import. Our data also demonstrates that nuclear compartmentalization by chromatin surface nucleation can be insensitive to the DNA-to-cytoplasm ratio. Thus, we speculate that cells will exploit these two complementary mechanisms to perform distinct functions. For example, surface nucleation may ensure that the necessary components for proper chromatin and nuclear functioning are partitioned into the nucleus regardless of cell size during the rapid divisions in early embryogenesis or asymmetric divisions. Competition for the import machinery may lead to component dilution during asymmetric divisions or early development, providing a differentiation or signaling response. Consistent with this, H1.8 is not titrated within the nucleus during early embryonic development^14^ while other proteins that are imported, such as H3, are titrated^4,6,8^. Additionally, surface nucleation may play a key role in mitotic bookkeeping and maintenance of cellular memory across multiple cell cycles.

Physical parameters of surface condensation, such as wetting angle and nucleation rate, can be tuned to provide a diverse range of functionalities^39–42^. The wetting angle determines whether droplets form on the surface—as our data shows— or whether the chromatin surface is completely sealed off by a liquid film. Droplet formation on the chromatin surface can lead to the specific formation of nuclear stores that can be readily used. Sealing the chromatin surface with a liquid film may explain the mechanical clustering of chromatin that leads to the non-specific exclusion of large particles from the cytoplasm (ki-67^43^), or could also play a key role in nuclear envelope formation (lap2^30^ and LEM^29^). The rate of condensation on chromatids could in turn be regulated by the properties of cytoplasmic condensation which can buffer its nucleation or make it sensitive to cell volume. As surface nucleation on chromatin provides a versatile mechanism for nuclear partitioning, it will be interesting to investigate the condensation behaviour of different nuclear components, and how that behaviour is regulated during the different stages of the cell cycle.

Altogether, our work highlights the significance of the physics of surface condensation and wetting as a key regulatory principle in intracellular biological systems, with implications beyond the self-assembly of structures like the nucleus.

## Extended Figures

**Extended Figure 1:**
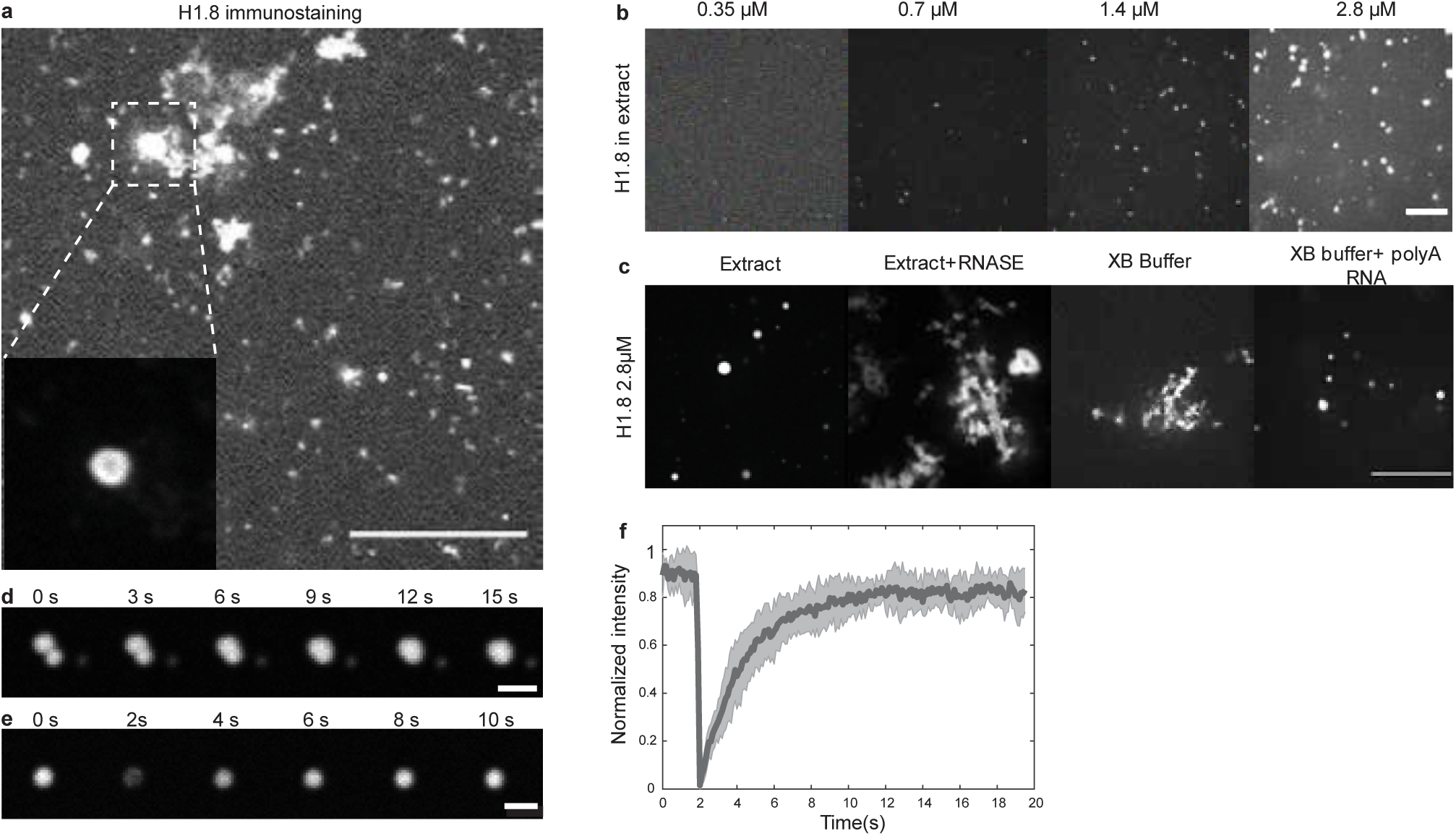
H1.8 undergoes liquid-liquid phase separation in egg extract. a, Immunofluorescence against H1.8 shows speckles and a few larger condensates in the cytoplasm. The inset shows a magnified view of a large condensate with different contrast to highlight a ring-like structure (see Extended Figure 2). Scale bar, 10 μm. b, Increasing the concentration of H1.8 in extract leads to an increasing amount of condensates indicating phase separation. Scale bar, 5 μm. c, RNA is necessary and sufficient for H1.8 to undergo phase separation. Removal of RNA from extract leads to protein aggregation in extract. The addition of H1.8 to buffer similar to extract salt concentration and pH leads to similar aggregation but are stabilized by addition of polyA RNA. Scale bar, 5 μm. d, H1.8 condensates in extract fuse upon persistent contact and undergo shape relaxation indicating liquid behavior. Scale bar, 2 μm. e, Representative images of fluorescent recovery upon photobleaching (FRAP) indicating quick molecular rearrangements typical of liquids. Scale bar, 2 μm. f, Quantification of FRAP of H1.8 condensates showing they recover most of their original intensity within 10 seconds (mean ± s.e.m, n=8).

**Extended Fig 2:**
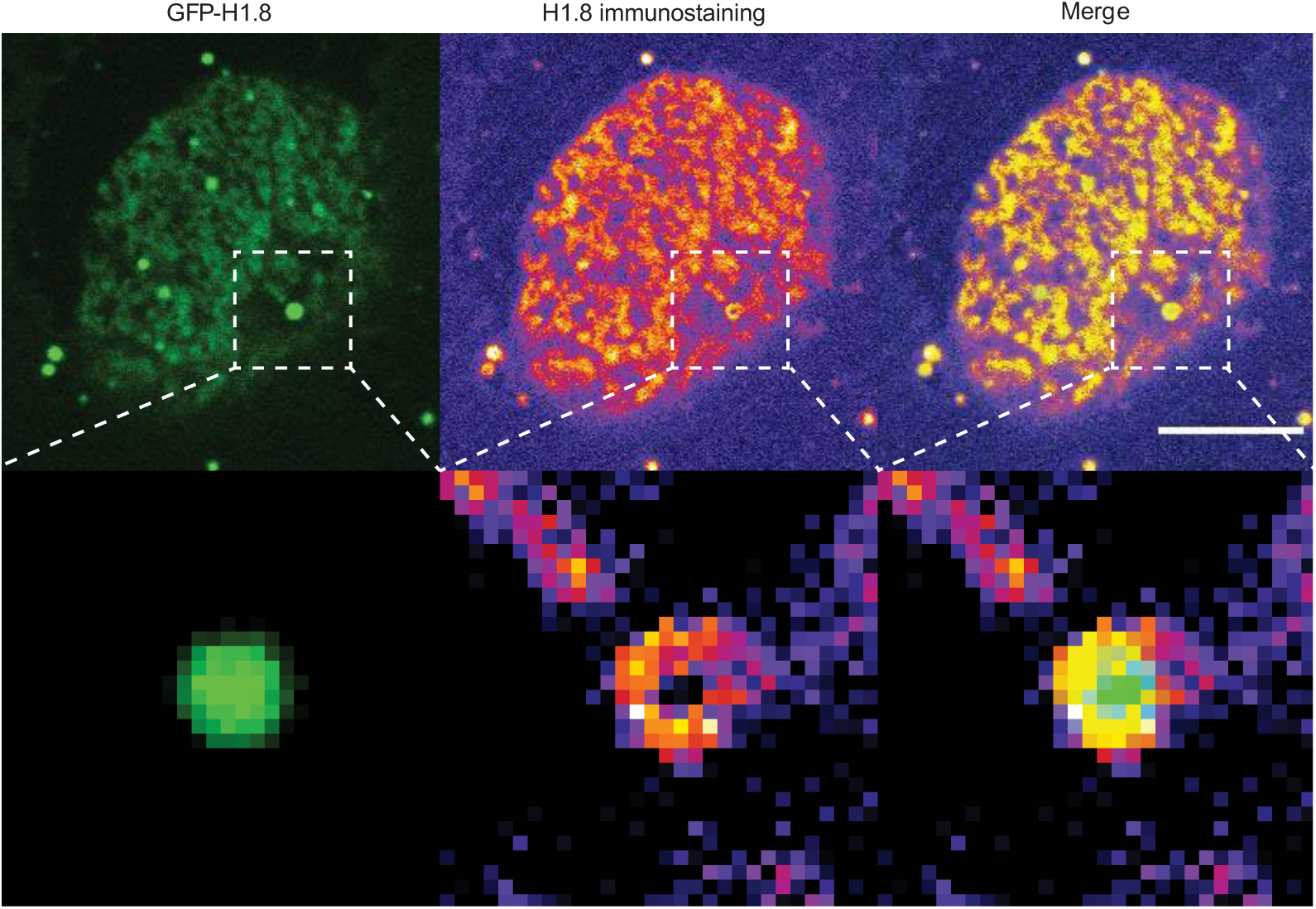
Immunofluorescence against GFP-H1.8 shows that the antibody only coats the surface of the condensates. The dotted box was magnified to highlight that the antibodies only surround the condensates. Due to this, smaller condensates can be easily missed when imaged through immunofluorescence. Scale bar 10 μm.

**Extended Fig 3:**
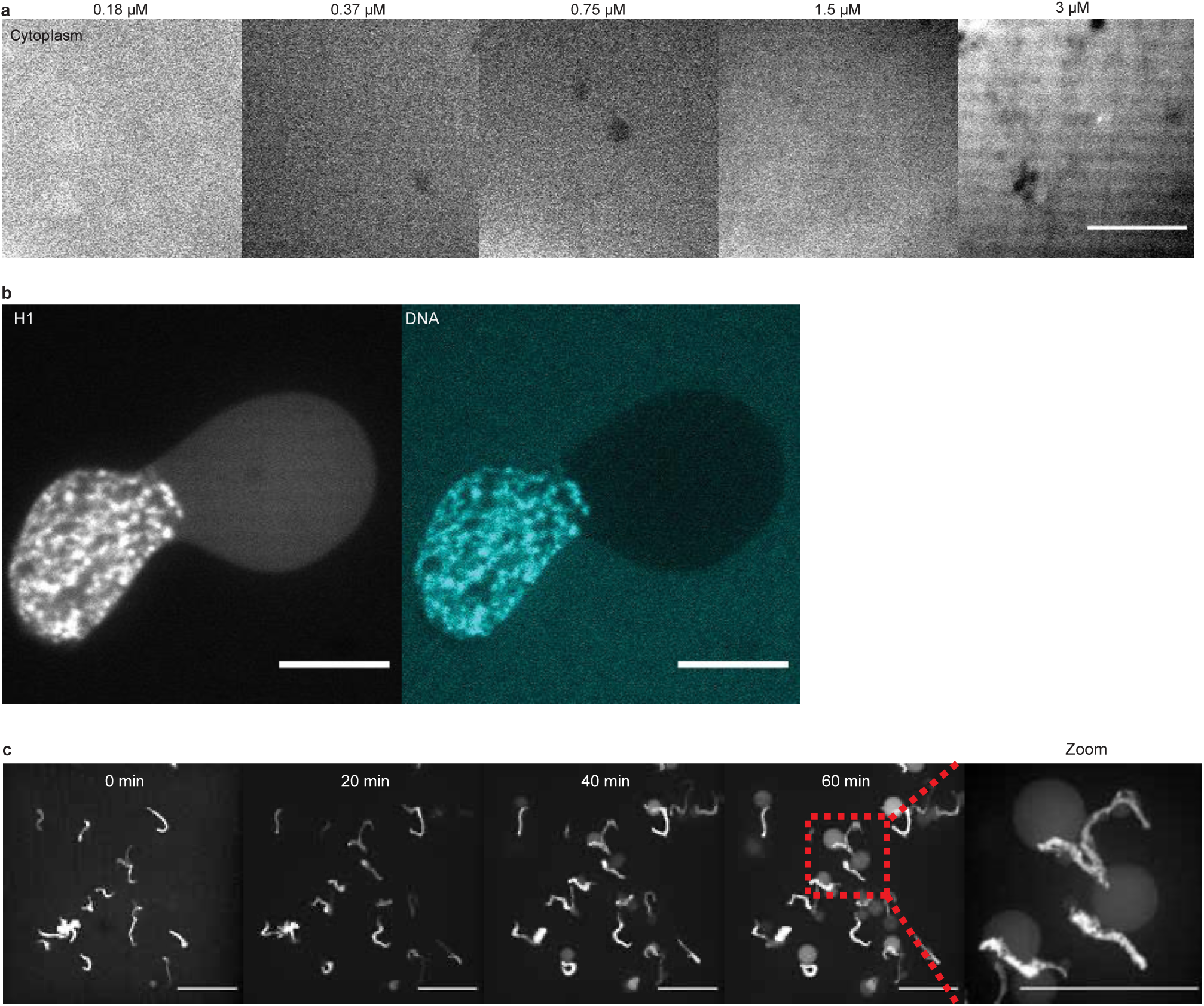
Somatic H1 doesn’t undergo phase separation but instead compacts chromatin. a, Somatic H1 (calf-thymus) remains diffusive in the cytoplasm across different concentrations. Scale bar, 10 μm. b, Somatic H1 leads to nuclear blebbing with condensed chromatin structures. Scale bar, 10 μm. c, Timelapse imaging shows that chromatin fails to expand in the the presence of somatic H1, but continued import leads to nuclear blebbing. Scale bar, 50 μm.

**Extended Fig 4:**
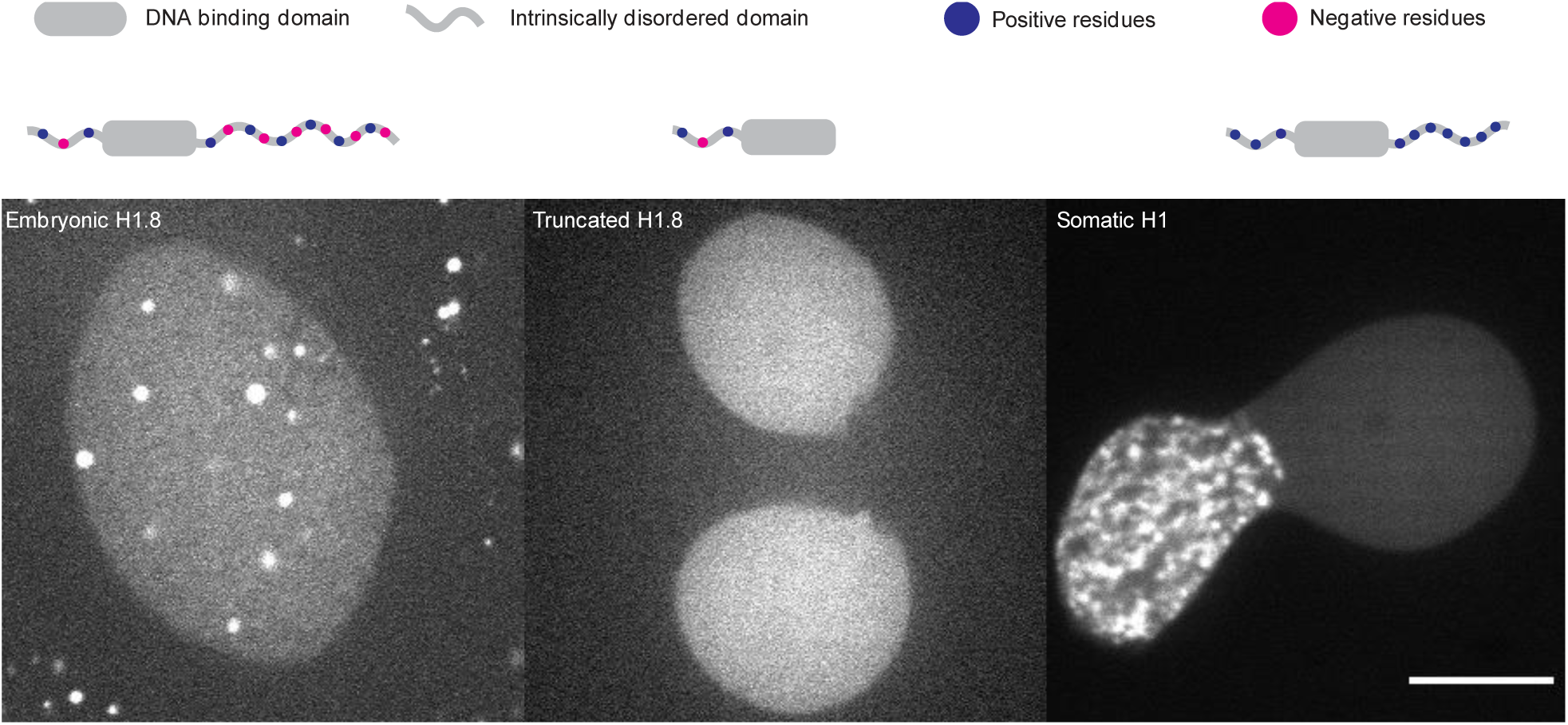
The charge distribution of IDRs drive condensation of H1 proteins. The IDR of embryonic H1.8 is zwitter ionic leading to formation of condensates in both the nucleus and the cytoplasm (left). Removing this IDR abolishes the formation of condensates (middle) while a highly positive IDR compacts chromatin within the nucleus.Scale bar, 10 μm.

**Extended Fig. 5:**
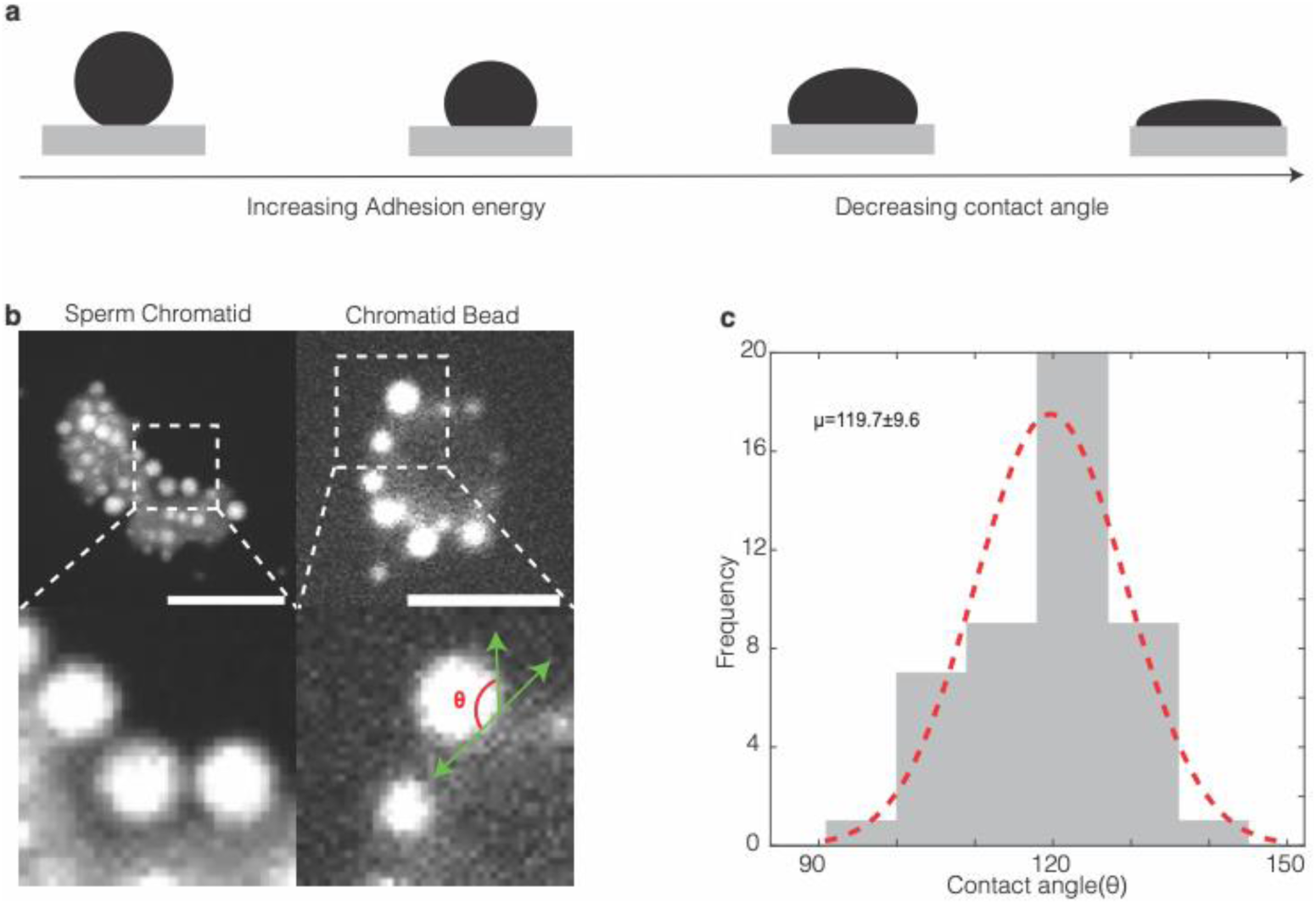
H1.8 partially wets the chromatin surface. a, Schematic representing how the contact angle changes with the adhesion energy. Low adhesion to the surface results in a high contact angle, while high adhesion energy leads to a low contact angle and complete wetting. b, H1.8 condensates on a chromatid (left) and DNA beads (right) show that they only partially wet the chromatin surface. The dotted box is magnified to show partial wetting more clearly. The contact angle is defined as shown. Scale bar, 5 μm. c, Histogram of contact angles from condensates (n = 47) on beads gives a mean ± s.d of 119.7 ± 9.6.

**Extended Fig. 6:**
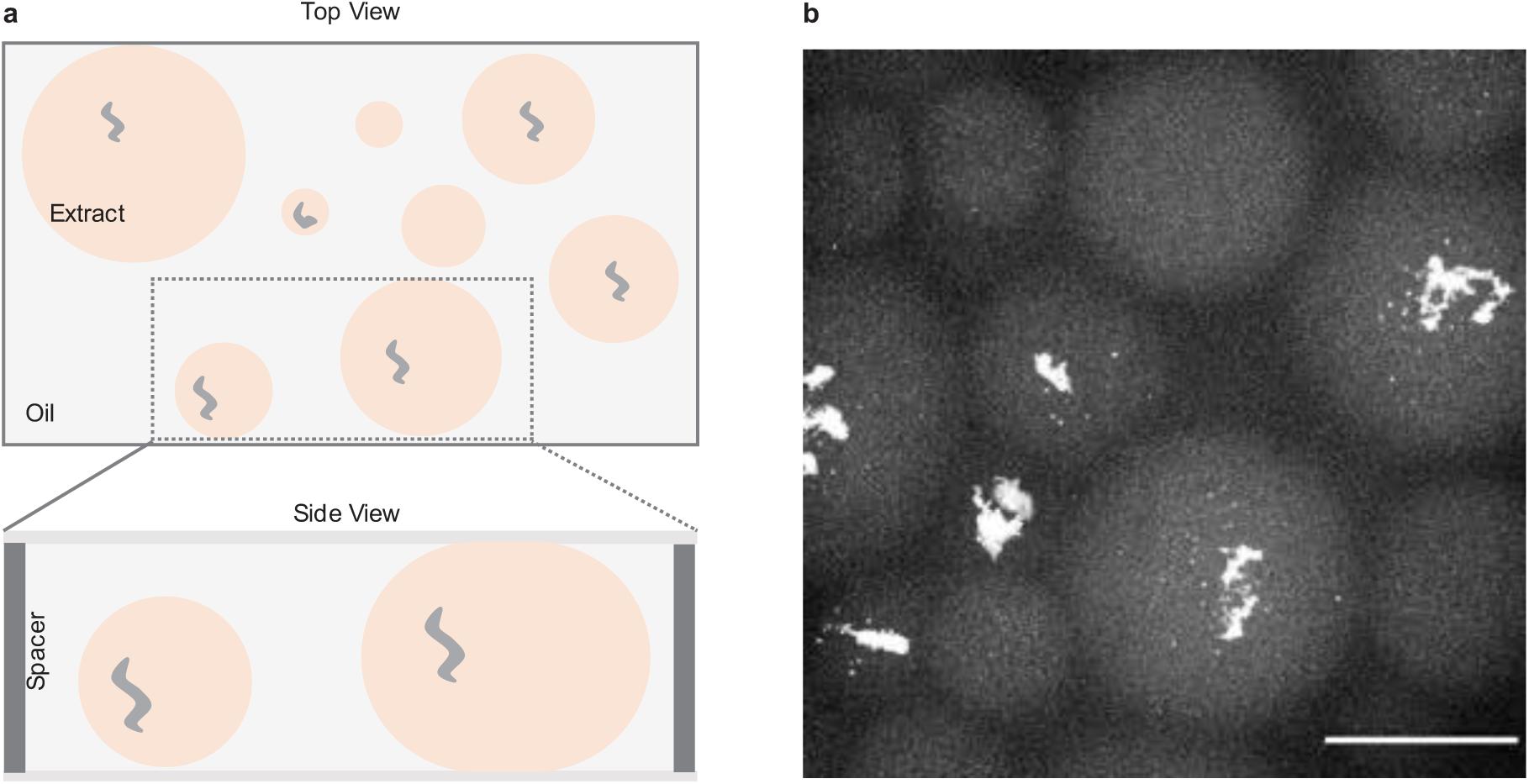
Chromatid assembly and H1.8 nucleation in cell-like compartments made by encapsulating *Xenopus laevis* egg extract using oil. a, Schematic of the imaging chamber used. Droplets are placed between a spacer to prevent compression. b, Representative images of chromatids in extract droplets of different volumes. Scale bar, 10 μm.

**Figure 11:**
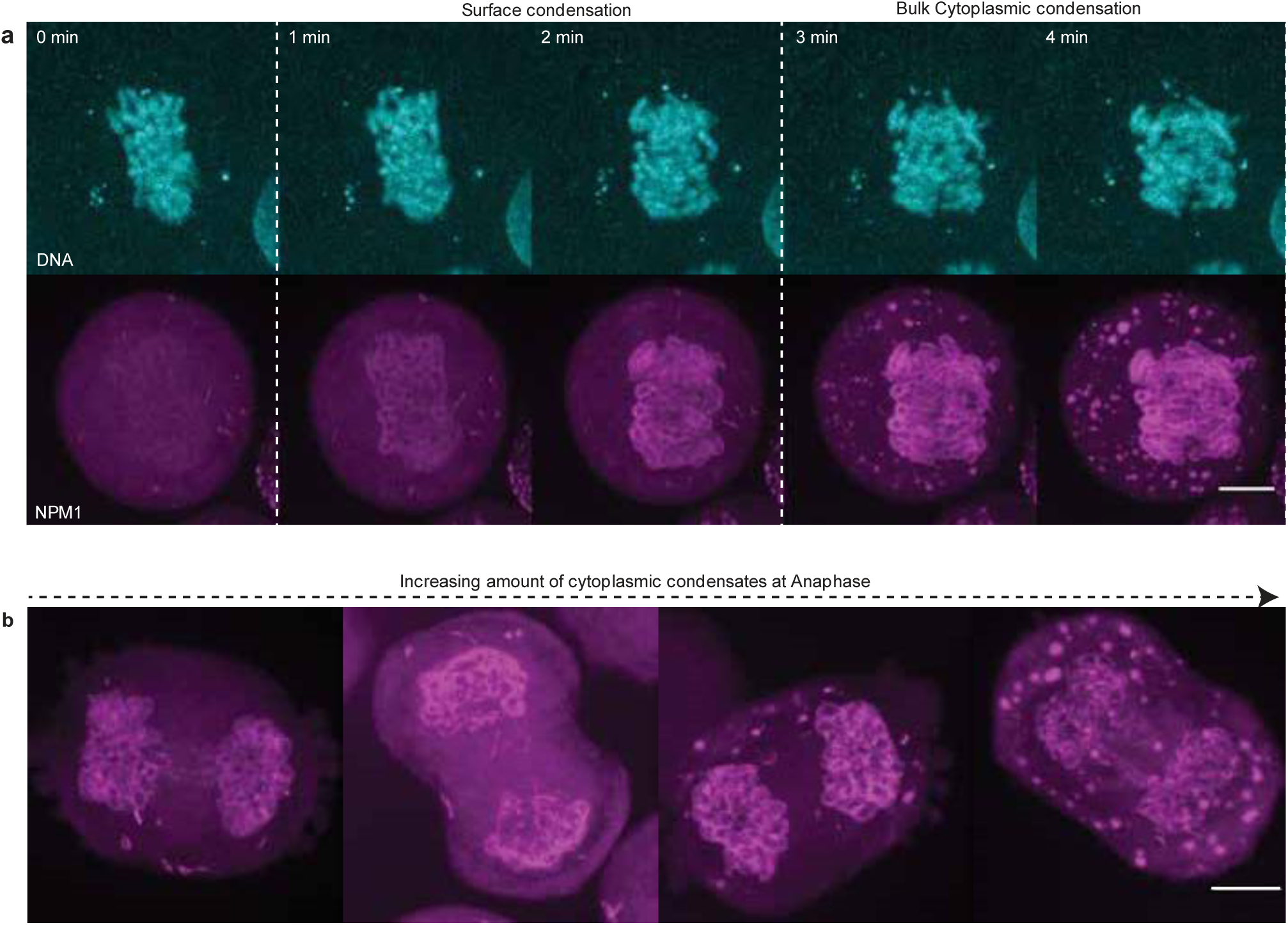
NPM1 forms condensates in the cytoplasm. a, A representative time-lapse image showing that NPM1 condenses on the surface of chromatids before forming cytoplasmic condensates. b, Increasing amount of cytoplasmic condensates in different cells (from left to right). There is considerable variability in the amount of cytoplasmic condensates between different cells in anaphase. All scale bar 10 μm.

## Acknowledgments

We would like to acknowledge M. Marass for his feedback on the manuscript. We would like to thank A. Bogdanova for discussions and help with protein purification as well as both the Protein Expression and Purification Facility and the Light Microscopy Facility at the Max Planck Institute of Molecular Cell Biology and Genetics. We would also like to thank the Funabiki lab for providing us with antibodies against H1.8. This work was supported by a DFG project BR 5411/1-1 (J.B., V.M.) and Volkswagen ‘Life’ grant number 96827 (J.B., V.M.), and a EMBO long-term fellowship (ALTF-1456-2015) (T.Q.).

## Author Contributions

V.M. and J.B. conceived the project. V.M. performed the experiments and image analsysis. J.B made the theoretical calculations. V.M. and T.Q. made the constructs for GFP-H1.8 and purified the protein. M.E. made the DNA beads. V.M. and J.B. wrote the manuscript.

## Author Information

**Cluster of Excellence Physics of Life, TU Dresden, Dresden, Germany**

Vasanthanarayan Murugesan, Maria Elsner & Jan Brugués.

**Max Planck Institute of Molecular Cell Biology and Genetics, Dresden, Germany**

Vasanthanarayan Murugesan, Maria Elsner & Jan Brugués.

**Max Planck Institute for the Physics of Complex Systems, Dresden, Germany**

Vasanthanarayan Murugesan, Maria Elsner & Jan Brugués.

**Center for Systems Biology Dresden, Dresden, Germany**

Vasanthanarayan Murugesan, Maria Elsner & Jan Brugués.

**European Molecular Biology Laboratory, Heidelberg, Germany**

Thomas Quail (Current affiliation)

## Supplementary Movie Captions

1) GFP-H1.8 nuclear condensates dissolve as the nucleus grows. Maximum intensity projections from 3D stacks of nucleus imaged through GFP-H1.8 shows that the condensates dissolve during nuclear growth, indicating slow import. The cytoplasmic background is removed for clarity. Scale bar, 10 μm. Relates to Fig 1e.

2) H1.8 condensate precedes nuclear formation. The video shows the transition of chromatids from metaphase to interphase. GFP-H1.8 (left) forms foci on the chromatid surface before the accumulation of NLS protein (right), indicating excess H1.8 enters the nucleus as condensates. Scale bar, 5 μm. Relates to Fig 2a.

3) Chromatin beads accelerate H1.8 condensation. The video shows the growth of GFP-H1.8 condensates in the cytoplasm (left) and on the surface of chromatinized beads (right). Scale bar, 5 μm. Relates to Fig 2e.

4) NPM1 condenses on the surface of chromatid surface to partition into the nucleus. The video shows the condensation of NPM1 on chromatid surface in anaphase and its subsequent incorporation and coarsening in late telophase. Scale bar, 10 μm. Relates to Fig 4a.

## Methods

### Xenopus laevis

*Xenopus laevis* adults were handled according to established protocols. The experiments were approved and licensed by the local animal ethics committee (Landesdirektion Sachsen, Germany; license no. DD24-5131/367/9) and carried out following the European Communities Council Directive 2010/63/EU on the protection of animals used for scientific purposes, as well as the German Animal Welfare Act.

### Reconstitution of nuclei/chromatids using Xenopus egg extracts

*Xenopus laevis* cycling egg extract capable was prepared as described previously in ref^44^. Briefly, unfertilized eggs were collected and dejellied using a 20 % (w/v) L-cysteine solution and were activated by adding 0.5 μg/mL calcium ionophore A23187, packed in a tube, and crushed by centrifugation. The cytoplasmic extract was then isolated and 10 μg/mL of a cockatil of protease inhibitors (LPC: Leupeptin, Pepstatin, Chymostatin) and Cytochalasin D were added. To reconstitute the nucleus and image protein localization, demembranated sperm were added (final concentration 300 to 1000 sperm/μL) along with recombinantly purified GST-mCherry-NLS (final concentration∼1 μM) and GFP-H1.8 (final concentration∼2.8 μM) and was kept on ice for ∼ 5 min. The reaction mixture was flicked and was gently pipetted onto a 35 mm imaging dish and spread with a pipette tip to form a thin layer, covered with mineral oil, and then imaged.

Cytostatic factor (CSF) arrested Xenopus laevis egg extract was prepared as described previously in ref^45^. It was prepared similar to cycling extract except instead of activating the eggs with calcium ionophere, they were incubated with a EGTA buffer (10 mM HEPES, pH 7.7, 50 mM sucrose, 100 mM KCl, 1 mM MgCl_2_, 0.1 mM CaCl_2_, 5 mM EGTA, 2 mM MgCl_2_) which chelated excess Ca^2+^ and keeps the extract in metaphase. To encapsulate extract in droplets, we added 15 μL of extract into 75 μL of Novec HFE-7500 oil supplemented with 5% (w/w) Pico-Surf surfactant (Sphere Fluidics) and flicked gently. We produced droplets with sizes between 10 – 100 μm in diameter. They were gently laid on a microscopic slide stuck with imaging spacer stickers of a 120 μm depth (70327-8S, Electron Microscopy Sciences). A coverslip was attached to the top of the spacer and then imaged.

To study to the effect of cytoplasmic condensates, GFP-H1.8 was added and incubated at 20 °C for one hour. Then, sperm was added and imaged.

### Protein Purification

We purified *X. laevis* embryonic linker histone H1.8 by modifying a published protocol from ref^20^. An in-house vector with H1.8 tagged with N-terminal eGFP tag and 6x Histidine-tag was transformed in T7 Express cells (enhanced BL21 derivative, NEB C2566I) and grown to an OD of ≈ 0.7 at 37 °C, whereupon 1 mM isopropylthio-β-galactoside was added to induce protein expression for 14-16 hours at 18 °C. All subsequent steps were carried out at 4 °C. Cells were collected and resuspended in lysis buffer (1× PBS, 20 mM imidazole, 500 mM NaCl, 2.5 mM MgCl2, 1 mM DTT, cOmplete EDTA free protease inhibitor cocktail, and Benzonase). The cells were lysed and centrifuged at 25,000 rpm for ∼ 30 min. The supernatant was collected and run through an immobilized metal ion affinity chromatography column. The protein was subsequently eluted with an elution buffer (1× PBS, 250mM imidazole, 500mM NaCl). The correct fractions were collected and dialyzed into PBS supplemented with 500mM NaCl, concentrated using Amicon Ultra centrifugal filters (10k cutoff), aliquoted, snap frozen using liquid nitrogen, and stored at -80 °C.ref purify GST-mCherry-NLS, a plasmid (a gift from Daniel Levy, University of Wyoming) containing an N-terminal GST-mCherry tag and a nuclear localization sequence (NLS) was transformed into T7 Express cells (enhanced BL21 derivative, NEB C2566I) and grown to an OD of 0.4-0.7 at 37 °C, whereupon 1mM isopropylthio-β-galactoside was added to induce protein expression for 14-16 H at 18°. The cells were resuspended in lysis buffer (1× PBS, 250 mM KCl, 2.5mM MgCl2, 1mM DTT, cOmplete EDTA free protease inhibitor cocktail; Roche, Benzonase), lysed, collected the supernatant, and ran over a GST-column. The protein was eluted with 50mM HEPES pH7.8, 10mM glutathione, and 10mM DTT, and dialyzed into XB buffer, spin-concentrated, snap-frozen, and stored at -80 °C.

### Immunofluorescence

The fixation and immunostaining were performed by using a previously published protocol from ref^21^. Briefly, 25 μL of extract, supplemented with sperm, was added to 2 ml of fixation buffer (80 mM K-PIPES pH 6.8, 1 mM MgCl2, 1 mM EGTA, 30% glycerol, 0.1% Triton X-100, 2% formaldehyde) and incubated for ∼ 5 min at RT, and layered onto 5-ml cushion (80 mM K-PIPES pH 6.8, 1 mM MgCl2, 1 mM EGTA, 50% glycerol) and spun onto a coverslip at 4500 rpm at 18 °C for 20 min. The coverslips were incubated overnight with antibody dilution buffer (TBS buffer supplemented with 2% BSA). Antibodies against *X. laevis* H1.8 (rabbit polyclonal, a gift from Funabiki lab, The Rockefeller University, New York) were added to the antibody dilution buffer to a final concentration of 4 μg/mL and incubated for 1 H at room temperature. The primary antibody was removed and the coverslip was washed three times with TBS buffer. Secondary antibodies against rabbit conjugated with Alexa Fluor 568 diluted to 4 μg/mL were added and incubated for 1 H at room temperature. Finally, the coverslip was incubated with siRHoechst or Hoechst (10 μg/mL) for 5 minutes. The coverslips were then washed with TBS buffer and mounted using 90% glycerol, and 10% PBS, sealed with nail polish, and stored at -20 °C.

### DNA beads preparation

25 μg of pTH303 plasmid was cut with BamH1 and NotH1, purified by ethanol precipitation, and their ends were biotinylated via a Klenow reaction supplemented with 25μM Biotin-ATP. Dynabeads protein A (catalog number 10001D, Thermofisher) was washed and then incubated with biotinylated DNA (∼10 μg of DNA for 50 μg of beads) overnight at room temperature under rotation. The beads were isolated using a magnet, washed, and then resuspended in XB buffer (10 mM HEPES, pH 7.7, 10 mM KCl, 1 mM MgCl_2_, 0.1 mM CaCl_2_). The beads were then isolated again and resuspended in extract and incubated in ice for ∼ 30 min for chromatinization. Then GFP-H1.8 was added to the extract with chromatinized beads to a final concentration of 2.8 μM and then imaged.

### Phase separation of H1.8 in extract/buffer

GFP-H1.8 or somatic H1 (H13188, ThermoFisher Scientific) was diluted in extract to obtain different concentrations (0.35 to 2.8 μM) and was incubated for 1 H at 20 °C before imaging. To remove RNA, RNase A (catalog number EN0531, Thermofisher Scientific) was added to the extract at a final concentration of 10 ng/ul and incubated on ice for 30 minutes, before adding GFP-H1.8 to a final concentration of 2.8 μM. For in vitro phase separation assays, GFP-H1.8 was added to XB buffer (10 mM HEPES, pH 7.7, 50 mM sucrose, 100 mM KCl, 1 mM MgCl2, 0.1 mM CaCl2). To test dependence on RNA, poly(A) RNA (Catalog number P9403, Sigma) was added to a final concentration of 100 ng/ul of buffer to XB buffer before adding GFP-H1.8.

### Fluorescence recovery after photobleaching (FRAP)

We performed FRAP using a femtosecond laser ablation setup was composed of a mode-locked Ti: Sapphire laser (Coherent Chameleon Vision II) oscillator coupled into the back port of the Nikon spinning disk microscope and delivering 140 fs pulses. Photobleaching was performed using a wavelength of 800nm and typically a power of 30 mW before the objective. GFP-H1.8 condensates of diameter greater than 2 μm, were partially bleached and the fluorescence recovery was recorded at a rate of 100 ms/frame for 20 seconds.

## Cell culture and drug treatment

HCT116 cell lines with native NPM1 tagged with mNeon green was a gift from the Zechner and Hyman lab. The cells were cultured at 37 ° with 5 % CO_2_ in McCoy medium (Gibco, cat # 366000) supplemented with 10% fetal bovine serum (Gibco, catalog # 26140-079) and 0.5 mg/ml of Penicillin-Streptomycin (Gibco, catalog # 10378-016). The imaging of cells from metaphase to interphase were performed using a slightly modified protocol from ref^33^. Briefly, cells were treated with 50 μM monastrol (Sigma-Aldrich, catalog # M8515), subjected to five medium washes, and dislodged from the flask surface by tapping the side of the bottle. Subsequently, cells were centrifuged at 500 g for 2 minutes, transferred to an 8-well slide (Ibidi, catalog # 80826), and incubated at 37 ° with 5 % CO_2_ in a media with siRHoechst (2 μg/mL) for 20 minutes before live imaging.

## Image acquisition and data analysis

### Image acquisition

The extract samples were usually kept on ice before imaging. The images were taken typically 5 to 10 minutes after the samples were transferred to the microscope. Nuclei and chromatids in extract were imaged using a Nikon spinning disk microscope (Ti Eclipse), an EMCCD camera (Andor iXon DU-888 or DU-897), a 20x 0.75 NA objective or a 60x 1.2 NA water immersion objective. GFP-H1.8 condensates were imaged using the 60x 1.2 NA water immersion objective. DNA beads were imaged using the same set-up but with a 100x 1.4 NA oil immersion objective. The AndorIQ software was used for image acquisition. The temperature in the room was maintained at 20 °C. HCT116 cells were imaged with Nikon Ti2 with Yokogawa CSU-W (NIS Elements 5.4), a Hamamatsu Orca Fusion BT and a 100x 1.4 NA water. All image analysis and segmentation was done using Fiji.

### Timelapse imaging of cycling extract

Cycling extract supplemented with sperm, GFP-H1.8, and mCherry-NLS were imaged using a spinning disk confocal microscope with a 20x 0.75 NA objective. Z-stacks were acquired with 1 μm optical sectioning every ∼ 2 min for 3 H. Then, the images were projected using their maximum intensity, and nuclei were segmented using the NLS signal using the Otsu method for histogram-based thresholding. Then using the analyse particles tool, the ROIs of the nuclei were identified. Based on the ROI, the mean intensity and the NLS and H1.8 was measured. The mean intensity was normalized from 0 to 1, and an average of all nuclei at a given time point was calculated. The beginning of the interphase was estimated by the appearance of the NLS signal and was assigned to timepoint zero.

### Measuring condensation on DNA beads

The DNA beads were added to extract with GFP-H1.8 and imaged using a spinning disk with 100x 1.4 NA oil immersion objective, with 0.5 μm optical sectioning every ∼ 2 min. The DNA beads were manually cropped and projected using their maximum intensity, and the condensates were segmented using the Minimum method for histogram-based thresholding in ImageJ. Then, we used the ‘Watershed’ function to ensure that the condensates were individualized. Then the ‘Analyze Particles’ function was used to measure the area of the condensates. The volume was estimated from the measured area. Similarly, the cytoplasmic area was cropped and the same analysis protocol was used.

### Measuring condensation on chromatids encapsulated in extract

CSF-extract supplemented with GFP-H1.8 and NLS was encapsulated in oil. The NLS channel was used to confirm that the extract was in metaphase. They were imaged using a 60x 1.2 NA water immersion objective with a 1 μm optical sectioning after 30 min to 40 min. The volume of condensates on chromatids was measured similarly to the DNA beads mentioned above. The volume of the encapsulated cytoplasm was estimated through the radius, which was measured manually.

### Scaling of condensate volume with cytoplasmic volume

We use a simple mass action calculation to fit the dependence of the volume of surface condensates with cytoplasmic volume in the absence of cytoplasmic condensates. Let M_t_ be the total mass of the protein encapsulated inside an extract droplet of volume V. From that mass, we denote M_b_ the fraction of mass that binds to DNA and M_s_ the soluble cytoplasmic fraction, so that ***M_t_*** = ***M_b_*** + ***_s_***. In this model, we consider the rate of protein binding to DNA as proportional to the cytoplasmic concentration C_s_, with a constant of proportionality α, 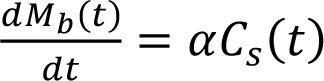. The soluble concentration in the cytoplasm is given by 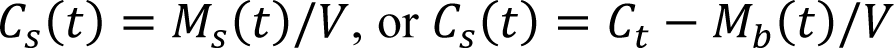, where *C_t_* is *M_t_*/*V* and is the initial concentration at time *t* = 0. Thus, the temporal evolution of the soluble concentration as condensates are nucleated on the chromatin surface is given by 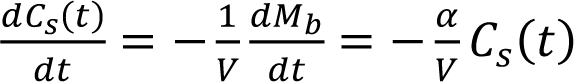, which leads to 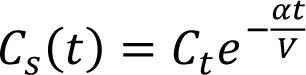. Substituing this solution for the soluble concentration into the equation describing the mass of condensates leads to, 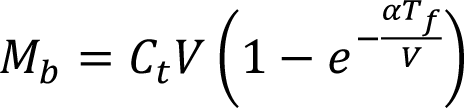, where *T_f_* is the duration of the experiment. This expression describes how the total mass of condensates bound to DNA change with the cytoplasmic volume. For a protein that phase separates in the cytoplasm, the soluble concentration will always be *C_sat_*. Substituting *C_s_* with *C_sat_* we get in this case *M_b_* = *C_sat_*α*T*.

### Measuring NPM1 localisation

The cells were imaged using a 100x 1.4 NA oil immersion objective with a 0.5 μm optical sectioning. The cells were live imaged for 20 minutes as they transitioned from metaphase into late telophase. The mean intensity of NPM1 on the chromatin surface and chromatin matrix plotted in Fig. 4c were measured manually. The chromatin surface and the matrix were identified using the DNA channel labelled with siRHoechst. Images in Fig 4a, b were captured at specific cell cycle stages using increased exposure and laser power settings to enhance image quality.

